# miR-21-5p and miR-30a-5p are identical in human and bovine, have similar isomiR distribution, and cannot be used to identify xenomiR uptake from cow milk

**DOI:** 10.1101/275834

**Authors:** Bastian Fromm, Juan Pablo Tosar, Yin Lu, Marc K. Halushka, Kenneth W. Witwer

## Abstract

microRNAs (miRNAs) are often highly conserved across species, but species-specific sequences are known. In addition, miRNA “isomiRs” arise from the same precursor molecule but differ in post-processing length and modification, usually at the 3’ end. A recently published feeding study reported the intriguing result that two bovine milk-specific miRNAs were taken up into human circulation after ingestion of bovine milk. Unfortunately, this interpretation is based on annotation errors in a public microRNA database. Reanalysis using databses including the MirGeneDB database reveals that the miRNAs in question, miR-21-5p and miR-30a-5p, arise from 100% identical 5’ precursor sequences in human and bovine, and the putative bovine-specific isomiRs appear to be depleted, not enriched, in bovine milk. Thus, enrichment of these isomiRs in human blood is inconsistent with uptake of xenomiRs and likely betrays endogenous miRNA regulation in response to diet or technical artifact.

---

It was recently reported (1) that two bovine microRNAs (miRNAs) were detected in human blood plasma after ingestion of 1 liter of 1%-fat milk. Unfortunately, the premise of a bovine-human difference in these miRNAs is based on annotation errors in a public microRNA database (2).

Mature miRNAs are formed through consecutive cleavage events of a primary miRNA transcript by the ribonucleases Drosha and Dicer (3,4). Each mature miRNA has consensus 5’ and 3’ cleavage sites. However, instead of a single sequence defining a miRNA, a collection of “isomiRs” exist, with different 3’ end lengths and modifications. Even the most abundant specific sequence of a given miRNA is, on average, only 45% of total reads (5). Additionally, for 204 miRNAs in the public miRBase repository used by Wang et al (2), the designated “canonical” sequence is different from the most abundant sequence.

Wang et al (1) make the unexpected claim that miR-21-5p and miR-30a-5p have unique sequences in bovine and human. Specifically, the bovine version of each is said to include two additional 3’ nucleotides (nt) not found on the human miRNA, presumably based on miRBase (Figure 1A). However, at the DNA level, the human and bovine sequences are identical through the longer purported bovine miRNA. Thus, the 3’ isomiR families of human and bovine cannot be distinguished by nt differences, as also supported by the actual read counts in miRBase (not shown).

**Figure 1.**
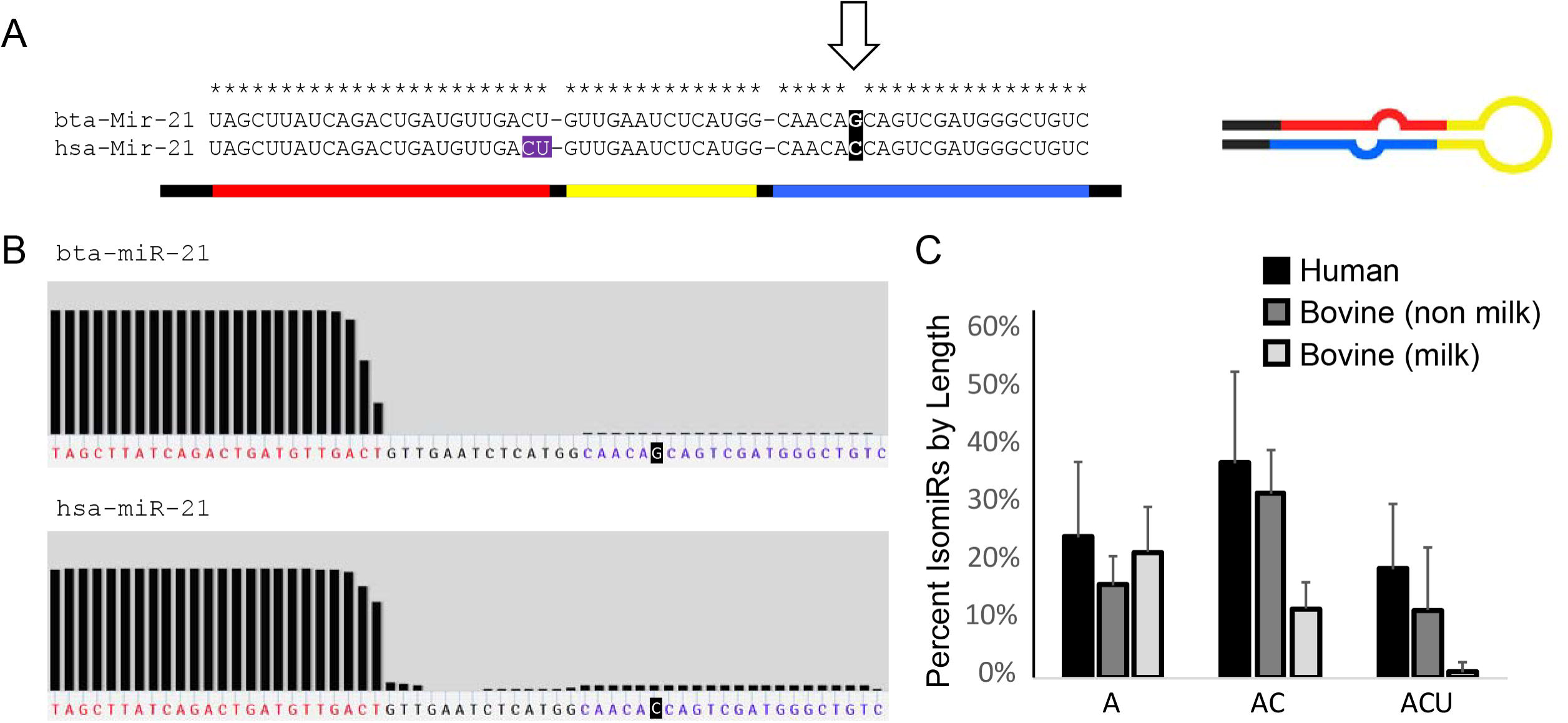
miR-21 in cow and human. A) bta- and hsa-miR-21-5p are 100% identical in bovine and human, (miRBase misannotation highlighted purple). Note the nt difference in the −3p arm, not studied by Wang et al. B) MirGeneDB isomiR coverage shows that human isomiRs are at least at the same length and level as in bovine. C) miR-21-5p isomiR distribution data from 122 human samples, 66 cow samples and 48 cow milk samples based on the canonical isomiR ending in A, AC or ACU. Other isomiRs are not shown.

Wang et al (1) would have been better served to view these miRNAs at MirGeneDB (6,7), an updated database. miR-21-5p and miR-30a-5p have the same mature miRNA sequence for both species (the longer, 24-nt sequence), which is shared across all vertebrates (not shown). MirGeneDB decay plots of miR-21-5p (Figure 1B) and miR-30a-5p (not shown) show the same 3’ cut site in bovine and human and suggest a high proportion of the 24-nt miR-21-5p with terminal ‘U/T’ in human.

Despite this clarification, it would still be possible that different isomiRs are favored in different species, akin to what we noted in cells (5). If there were extreme skewing towards a 22-nt miR-21-5p or miR-30a-5p in humans or a 24-nt version in cow milk, the data in this report would be intriguing. However, that is not the case. From small RNA-seq data processed in miRge (8), we obtained the miR-21-5p isomiR spectrum from 122 human samples (6 studies) and 114 bovine samples (4 studies), of which 48 were from milk. Reported data are limited to samples with >1,000 reads of the specific miRNA (data available upon request). As seen in Figure 1C, cow milk appears to be depleted, not enriched, in the 24-nt, pan-vertebrate consensus isomiR of miR-21. This apparent 3’ decay may occur in milk-secreting cells, in the biofluid, during industrial processing of milk, or a combination of the above. Transfer of milk isomiRs into human blood, if detectable, would tend to dilute, not increase, levels of 24-nt isomiRs in human circulation after milk intake. These data are counter to the proposal of Wang et al (1) that milk ingestion specifically increased the 24-nt isomiRs (their proposed bovine-exclusive miRNAs). Data for miR-30a are similar, with the highest values for the full-length isomiR being in human, not cow milk (not shown).

It is not clear why 24-nt isomiRs would appear to increase slightly after a milk meal. We suspect either technical factors or perhaps real biology of human miRNAs changing their levels during the day (9) or in response to a food bolus (10). Concerning technical issues, rhPCR (11) was designed to distinguish single nucleotide polymorphisms (SNPs), but in the application by Wang et al, no true SNPs exist. Perhaps this method could be applied to molecules with internal nt differences, such as the passenger strand (miR-21-3p) that was not studied (Figure 1A), but necessary assay development and controls would need to be done (11). We also note that if the longer (“bovine”) versions of these miRNAs were truly unique to bovine or cow milk, they should not have been present in humans at time 0; yet they were detected.

In conclusion, Wang et al (1) performed a study of dietary miRNA uptake based on annotation mistakes in one public database (2) and failed to take into account the distribution of reads of two isomiR families. Public data show 100% sequence identity for the two miRNAs, and cow milk is depleted, not enriched, in putative milk-specific isomiRs. This manuscript does not further inform us of whether or not xenomiRs enter mammalian circulation, but rather adds to the questionable science advancing that narrative (12).

## Acknowledgements

All authors analyzed data. MKH and KWW wrote the paper and had primary responsibility for final content. All authors read and approved the final manuscript.

